# New insight into the RNA-chaperon activity of nucleobindin 1

**DOI:** 10.64898/2026.05.22.727093

**Authors:** O.S. Kostareva, I.A. Eliseeva, A.I. Buyan, D.N. Lyabin, S.V. Tishchenko, A.O. Mikhaylina

## Abstract

Nucleobindin 1 (NUCB1) is a multifunctional conserved protein located in Golgi luminal, nucleus, extracellular and cytosolic pools. NUCB1 is multidomain protein comprised of a signal peptide, a DNA-binding domain, a leucine zipper and Ca^2+^ -binding domain. The multiple domains and localization of NUCB1 potentiates its interactions with various partners, such as DNA, Gαi3 protein, cyclooxygenase 2, LRP10 and RNA suggests its importance in the regulation of many cellular events. We revealed that NUCB1 contains three RNA-binding regions and able to interact with two RNA fragments. It was suggested possible variants of the participation of NUCB1 in the interaction of the two partially complementary RNAs. The RNA-binding properties of the NUCB1 were also confirmed *in vivo* experiments.

## 1. INTRODUCTION

Nucleobindin 1 (NUCB1, calnuc) is a conserved Ca^2+^ -binding eukaryotic protein formed homodimer and involved in many processes, such as the stress response, immune response, and calcium homeostasis. NUCB1 was first found in the B lymphocyte cell line from systemic lupus erythematosus (SLE)-prone mice (Kanai *et al*., 1992). NUCB1 has a signal peptide and penetrates in various compartments of the cell and in the extracellular space (Kanai *et al*., 1992). Expression levels of NUCB1 have been observed to be elevated in tumors of the stomach, lymphocytes, and colon, its secretion has been reported in AtT20 pituitary tumor cells (Vignesh *et al*., 2021). The importance of NUCB1 in tumorigenesis and autoimmune diseases (Mikhailina *et al*., 2024) also emanates from the fact that it expressed ubiquitously and implicated in several physiological activities, including cell signaling, stress response, receptor sorting, inflammation, and apoptosis.

NUCB1 is involved in the regulation of many processes via interactions with Gαi3 protein (Lin *et al*., 2000), carboxyl terminus of Hsc70-interacting protein (Xue *et al*., 2016), APP (Lin *et al*., 2007), lipoprotein LRP10 (Brodeur *et al*., 2009), cyclooxygenase 2 (Leclerc *et al*., 2008), the metalloproteinase MMP2 (Pacheco-Fernandez *et al*., 2020). Besides, NUCB1 plays a role of chaperone-like protein by interacting with the partially folded intermediates of amyloidogenic proteins (Kanuru, Aradhyam, 2017). Moreover, the protein binds to the enhancer box (E-box) DNA sequence of the Cripto-1 gene promoter, triggering cell epithelial–mesenchymal transition (Sinha *et al*., 2019).

However, despite the large amount of data on the partner molecules of NUCB1 its precise physiological and cellular functions poorly understood.

Recently we demonstrated the RNA-binding and RNA-chaperone activity of NUCB1 and shown that NUCB1 prefers single-stranded G-rich RNA sequence motifs (Mikhaylina *et al*., 2023). Many RNA-binding proteins can efficiently promote intra strand RNA annealing in vitro (Hentze *et al*., 2018). NUCB1 was detected in exosomes of cancer cells (Vignesh *et al*., 2021), indicating a potential expansion of the protein’s physiological and cellular functions via its interaction with mRNA and microRNA and/or modulating the mRNA/microRNA interaction.

It is known that prokaryotic (Quendera *et al*., 2020; Woodson *et al*., 20218) and eukaryotic proteins (Serin *et al*., 1997; Barraud *et al*., 2013; Feng *et al*., 2022) with RNA-chaperone activity typically possess multiple RNA-binding domains.

NUCB1 consists of several functional domains: a signal peptide, N-terminal DNA-binding domain (DBD), two Ca^2+-^binding EF-hand motifs (CaBD) and C-terminal leucine zipper (LZ). In this study, we showed that each of the three domains of NUCB1 (DBD, CaBD, LZ) interacted with NUCB1-binding RNAs (guanine containing single-stranded (ss) RNA). Based on the “molecular beacon” method, two possible variants of the NUCB1 involvement in the interaction of two partially complementary RNAs was suggested. Using high-throughput sequencing, the NUCB1 mRNA targets in the HEK293T human cell line searched, identified, and analyzed.

## 2. RESULTS

We showed (Mikhaylina *et al*., 2023) that NUCB1 binds to ssRNA region containing guanine (mandatory), as well as adenine and/or uridine. The oligocytidine RNA did not bind to NUCB1. Using molecular beacon method, we have been demonstrated that full-length NUCB1 is capable of unwinding an RNA hairpin, the loop of which contains the NUCB1 binding site.

The three truncated forms of the protein: CaBD, ΔN, ΔZ (Fig.1) interacted with specific RNA fragments with high affinity (Mikhaylina *et al*., 2023) and ΔN and ΔZ forms contained calcium-binding domain. It demonstrated that the CaBD is sufficient for the RNA-binding activity of the NUCB1, but it was unclear how necessary it is.

**Figure 1.**
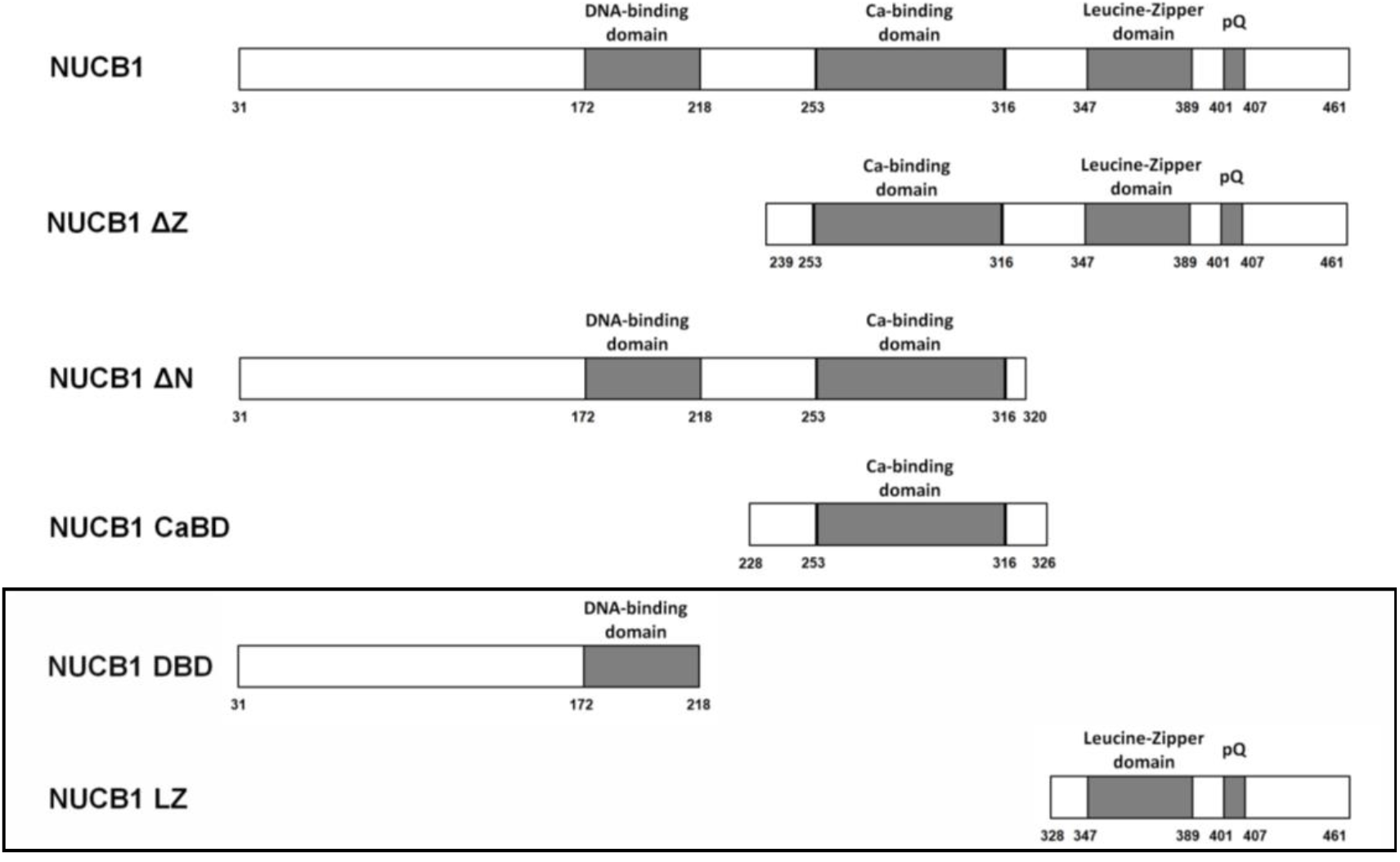
Schematic representation of NUCB1 and its truncated forms. The frame shown variants, which constructed in this study (NUCB1 DBD and NUCB1 LZ).

### 2.1. NUCB1 has three potential RNA-binding sites

In this work, we have obtained two truncated forms (Fig. 1) of the NUCB1 devoid of the calcium-binding domain: the DNA-binding domain (DBD, monomer) and the LZ domain (LZ, dimer). The RNA-binding activity of the NUCB1 and its truncated forms studied by the SPR method using as RNA fragment miR-200a-3p. (Table 1)

**Table 1.** Kinetic analysis of the interactions of NUCB1 and its truncated variants with miR-200a-3p.

| | Ka ($M^{-1} s^{-1}$) | Kd ($s^{-1}$) | K<sub>D</sub> (nM) |
| --- | --- | --- | --- |
| NUCB1 | $1.5 \times 10^4$ | $8.8 \times 10^{-5}$ | $5.8 \pm 0.3$ |
| DBD | $7.0 \times 10^3$ | $3.6 \times 10^{-4}$ | $51 \pm 7$ |
| LZ | $1.6 \times 10^4$ | $2.6 \times 10^{-3}$ | $162 \pm 10$ |
| CaBD* | $0.6 \times 10^4$ | $0.3 \times 10^{-3}$ | $5.2 \pm 0.2$ |

\*The data is taken from Mikhaylina *et al.*, 2023.

It has been demonstrated that NUCB1 truncated forms without CaBD have RNA-binding activity also. Thus, the calcium-binding domain is sufficient for the interaction of NUCB1 with RNA, but is not necessary. DBD and LZ domains of NUCB1 have RNA-bonding sites also.

### 2.2. RNA-chaperone activity of NUCB1, realized in the presence of two RNA fragments

The presence of multiple RNA-binding sites of NUCB1, demonstrated in this study, and experiments indicating its involvement in autoimmune (Mikhailina *et al*., 2024) and oncological processes (Vignesh *et al*., 2021), suggest a regulatory role for NUCB1 not only because of its interactions with proteins and E-box DNA. We assumed the participation of NUCB1 in interactions with two RNAs (f.e. mRNA and microRNA) *in vivo*. A question arose: “Can the chaperone activity of a NUCB1 be realized in the presence of two RNAs, each containing sites for potential interaction with the protein and with a target RNA?”

The miR-27b-3p microRNA pair and the predicted target (“seed”)sequence 3’UTR-FOXO1 were chosen as a model. The FOXO1 transcription factor orchestrates the regulation of genes involved in the apoptotic response, systemic lupus erythematosus, cancers, cell cycle, and cellular metabolism (Alikhani *et al*., 2005; Alikhani *et al*., 2010; Xing *et al*., 2018). FOXO1 expression is regulated by multiple microRNAs including miR-27b-3p (Guttilla *et al*., 2009; Duwe *et al*., 2023; Wang *et al*., 2019; D’Onofrio *et al*., 2023). The hsa-miR-27b-3p targets the 3′untranslated regions of FOXO1 mRNA to downregulate its expression, blunting the activation of the Akt/FOXO1 pathway. It was shown the interaction of conservative 426–432 region of FOXO1-mRNA 3′-UTR (Fig.2) with hsa-miR-27b-3p (D’Onofrio *et al*., 2023).

**Figure 2.**
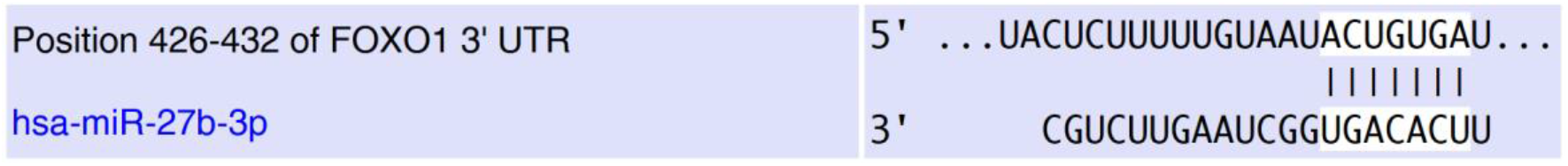
Alignment of hsa-miR-27b-3p with conserved region 426–432 of FOXO1 3′-UTR from miRNA-mRNA integration analysis using microRNA target prediction tools, as TargetScan (Agarwal *et al*., 2015). Predicted consequential pairing of “seed” region (top) and microRNA (bottom).

#### 2.2.1. NUCB1 can compete for the RNA binding site with another partially complementary RNA

We used the SPR method for investigation of NUCB1 binding to FOXO1 3’ UTR mRNA (426-432 nt) and miR-27b-3p fragments. The FOXO1 RNA fragment contained two guanines in loop (Fig.4a). It was shown that NUCB1 forms stable complex with FOXO1 RNA (Tabl.2). However, NUCB1 did not bind to fragment containing miR 27b-3p (Fig.4a). Apparently, a variant of the secondary structure of miR 27b-3p RNA realized in which guanines located not in the single-stranded loop, but in the double-stranded part, along with this the part of mRNA binding site (5’-terminal) was free for RNA-RNA interactions (Fig.4a).

The molecular beacon experiments were carried out as described in our work (Mikhaylina *et al*., 2023). The molecular beacon is a short RNA hairpin with a fluorophore at one end and a quencher at the other end. In the hairpin state ends of the beacon are brought together, and the fluorescence is quenched. When the RNA chaperone interacts with the molecular beacon’s loop nucleotides, the hairpin melts and fluorescence is restored.

In our experiment (assay 1), the molecular beacon included the sequence UAAUACUGUGA of 3’-UTR FOXO1 mRNA in loop of hairpin (fig.3a) with FAM (fluorophore) at the 5’-end and RTQ (quencher) at the 3’-end of the RNA hairpin (MB-FOXO). In the result, the beacon loop contained both microRNA- and NUCB1-binding sites; miR-27b-3p included the partially accessible mRNA-binding site only (Fig.3a). Upon addition of NUCB1 to the MB-FOXO in a 1:10 molar ratio, fluorescence significantly increased indicating that the RNA hairpin was melting. Adding a partially complementary miR-27b-3p to the RNA beacon practically did not change fluorescence, as did subsequent addition of the NUCB1 (Fig.3b). The investigation of the dynamics of RNA beacon melting by NUCB1 in the presence of partially complementary miR-27b-3p showed a rather sharp slowdown in the melting of beacon when microRNA was added (Fig. 3c). Apparently, partially complementary miR-27b-3p competed for the binding site on the MB-FOXO with the NUCB1.

**Figure 3.**
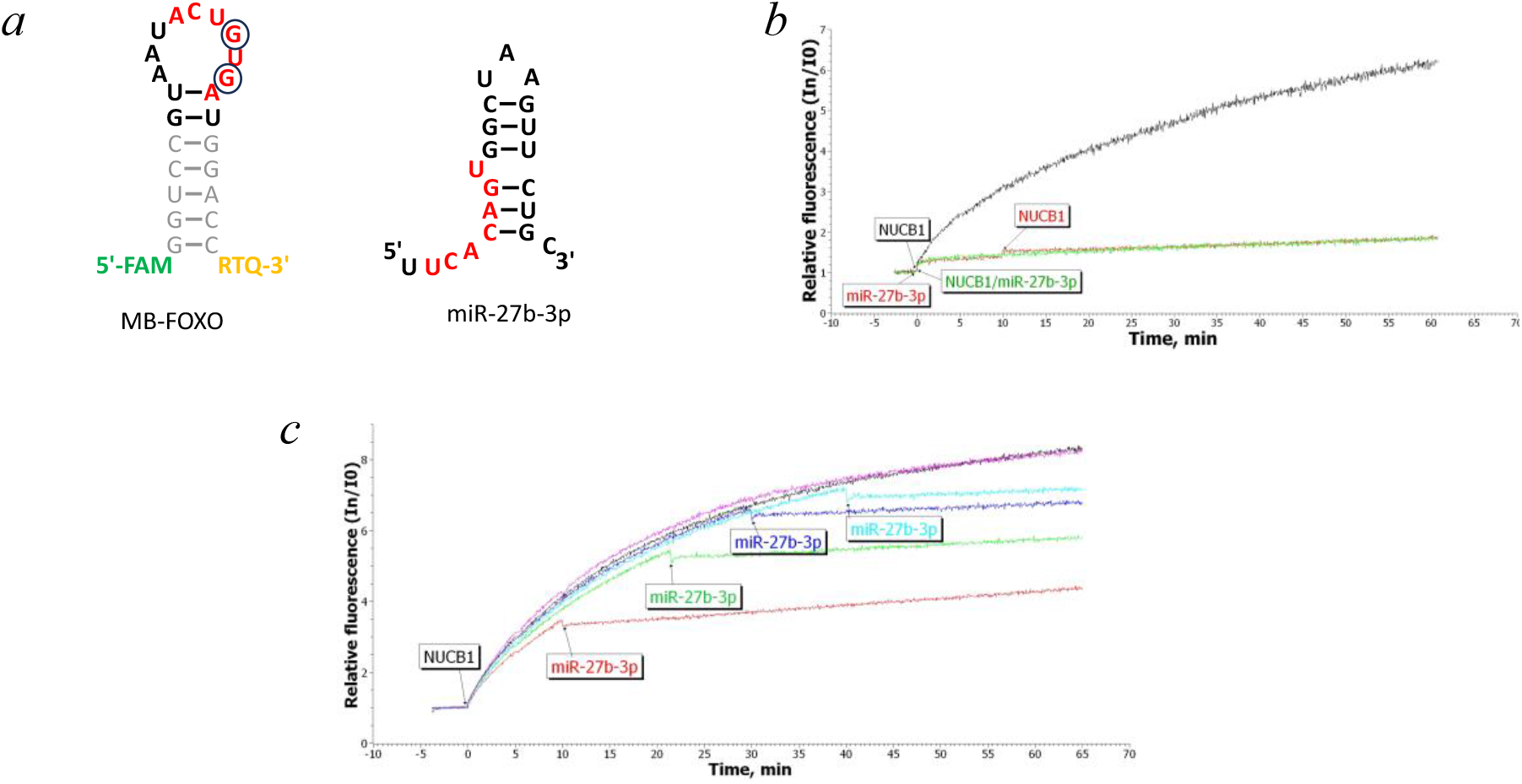
Molecular beacon melting (assay 1).

Secondary structures (a) of MB-FOXO and miR-27b-3p predicted using RNAfold web server. The complementary RNA regions highlighted in red.

Melting curves of MB-FOXO (b) after addition of NUCB1 (grey), after additional miR-27b-3p and added NUCB1 after 10 min (red), added mix NUCB1 and miR-27b-3p in molar ratio 1:1 (green). Melting curves of MB-FOXO (c) after addition of NUCB1 (black) and added miR-27b-3p after 10 min (red), 20 min (green), 30 min (violet), 40 min (blue), or water after 10 min (purple).

#### 2.2.2. NUCB1 can promote the interaction of two partially complementary RNAs if it interacts with both RNAs

In the following experiment (assay 2), we used new variant of mir-27b-3p (mir-27b-3p.1) containing, in addition to the mir-27b-3p sequence, a beacon helix sequence to form a single-stranded G-containing site in loop. The RNA-RNA interaction site can be located in an RNA loop 2 (Fig.4a).

**Figure 4.**
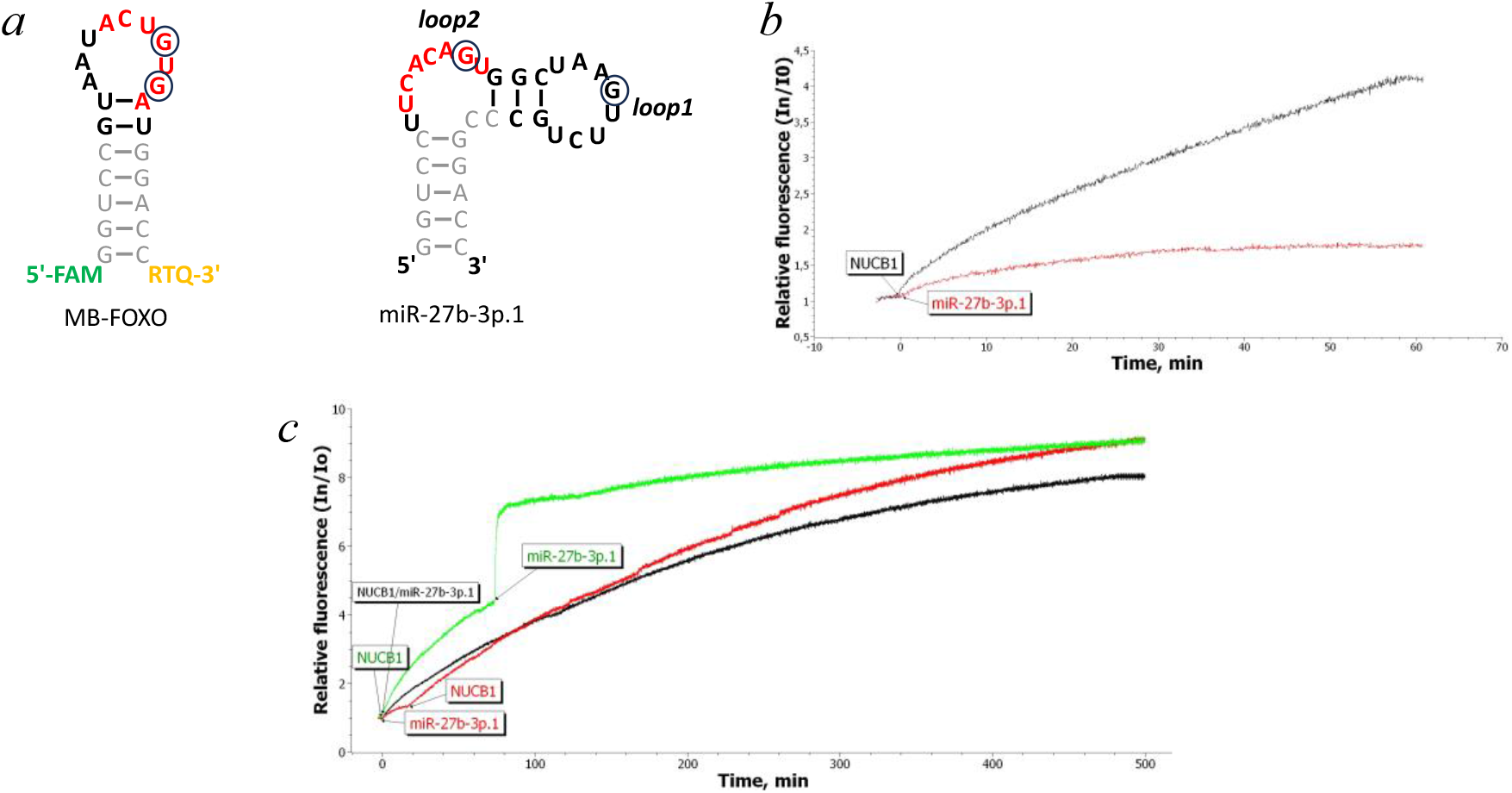
Molecular beacon melting (assay 2).

We have shown using the SPR method that NUCB1 forms stable complex with new variant of RNA fragment containing mir-27b-3p.1 (Table 2). The experiment with the MB-FOXO melting by NUCB1 repeated using mir-27b-3p.1 (fig.4b, c).

**Table 2.** Kinetic analysis of the interactions of NUCB1 with RNAs.

| RNAs | $K_a$ ($M^{-1} s^{-1}$) | $K_d$ ($s^{-1}$) | $K_D$ (nM) |
| --- | --- | --- | --- |
| FOXO1 | $9.9 \times 10^4$ | $3.6 \times 10^{-4}$ | $3.6 \pm 0.1$ |
| miR-27b-3p | No binding | No binding | No binding |
| miR-27b-3p.1 | $3.5 \times 10^5$ | $3.9 \times 10^{-2}$ | $111 \pm 10$ |
| SYDE1 | $2.3 \times 10^5$ | $7.1 \times 10^{-2}$ | $308 \pm 21$ |

In this case, NUCB1 melted MB-FOXO both in the absence (fig.4b) and in the presence of mir-27b-3p.1 (Fig.4c). The RNA beacon unwinding proceeds almost with the same intensity as in the absence of the mir-27b-3p.1. It can indicate the ability of the NUCB1 to bind to two RNA fragments at the same time.

Secondary structures of MB-FOXO and miR-27b-3p.1 (a). The complementary regions on both RNAs highlighted in red.

Melting curves of MB-FOXO (b) after addition of NUCB1 (black) or miR-27b-3p.1 (red).

Melting curves of MB-FOXO (c) by NUCB1 and adding after 40 min miR27b-3p.1 (green), adding NUCB1 in complex with miR-27b-3p.1 (black); adding miR-27b-3p.1 and NUCB1 after 10 min (red).

When mir-27b-3p.1 fragment added to the reaction mixture during the beacon unwinding process by NUCB1 (Fig.4c, green curve), a reproducible sharp increase in fluorescence intensity observed, followed by a slight further increase in intensity. The results of this experiment may indicate that the NUCB1 is capable of interacting with two RNAs containing a specific NUCB1-binding site at the same time and promote RNAs interaction with each other.

The next experiment (assay 3) was set up for further research of this phenomenon. We used mir-27b-3p.1 as beacon (MB-mir) and a FOXO1 3’-UTR mRNA fragment (FOXO1) as a competitive RNA (fig. 5a).

**Figure 5.**
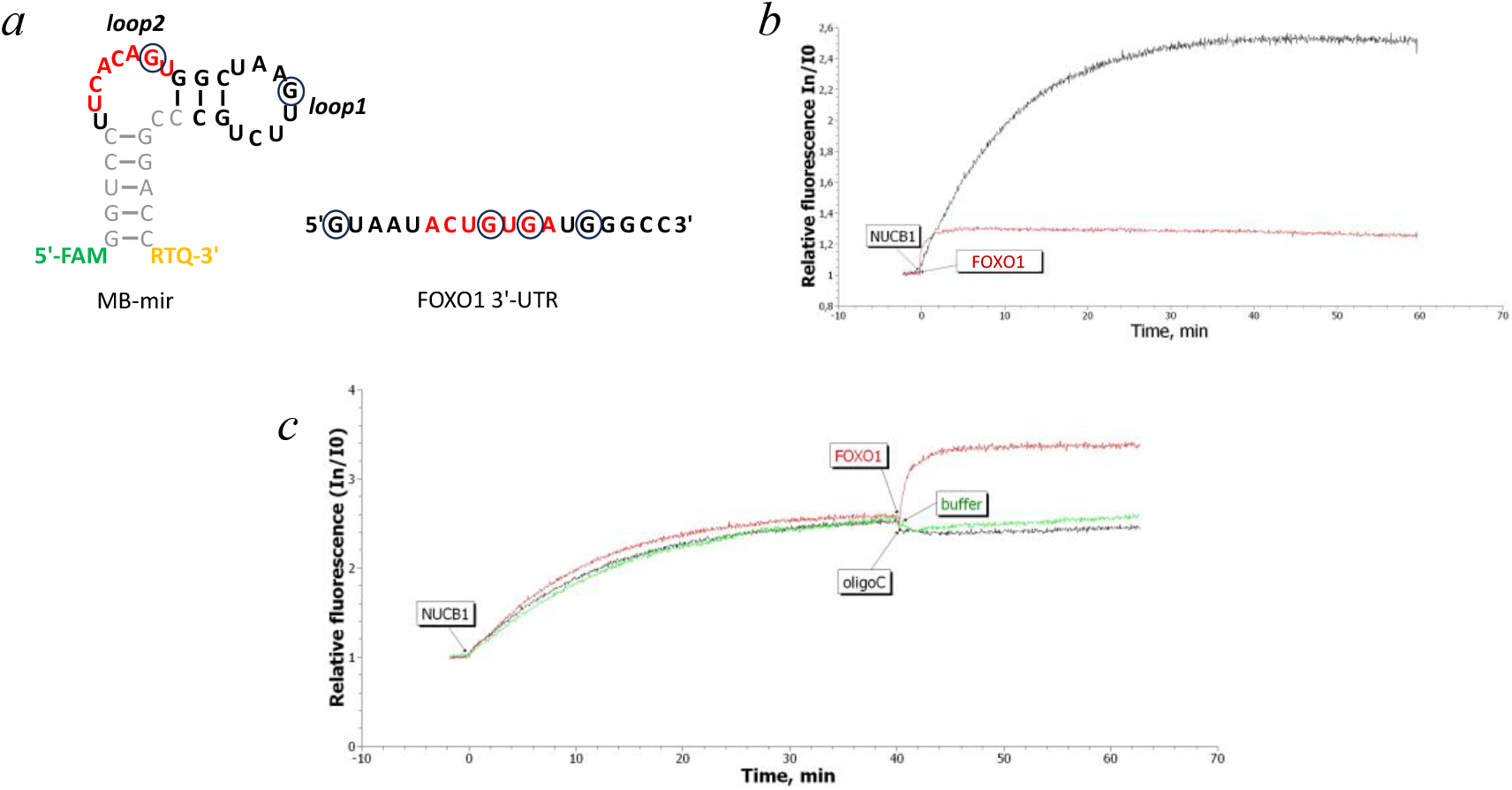
Molecular beacon melting (assay 3). The complementary regions on both RNAs highlighted in red.

Secondary structures of MB-mir and FOXO1 (a).

Melting curves (b) of MB-mir after addition of NUCB1 (gray) or FOXO1 (red).

Melting curves of MB-mir by NUCB1 (c) and adding at 40 min buffer A (green) or oligo(C)RNA (black) or FOXO1 (red).

Adding FOXO1 during beacon melting by NUCB1 increased in fluorescence also (fig. 5c, red curve), which may indicate that FOXO1 fragment interacts not only with NUCB1 but with RNA binding site on MB-mir also. It is worth noting that adding oligoC, with which neither RNA beacon nor NUCB1 not bind to (Mikhaylina *et al*., 2023), had no effect on beacon melting (Fig.5c, black curve).

### 2.3. The searching of NUCB1 mRNA targets in vivo

HEK-293T cells were transfected with a plasmid carrying the *NUCB1* gene, and mRNAs associated with NUCB1 were purified by immunoprecipitation (IP) followed by RNA sequencing. To identify mRNAs specifically associated with NUCB1, we compared the IP samples to total RNA and IgG controls. As shown in the volcano plots (Fig. 6), NUCB1 associates with RNA in HEK-293T cells, although the number of specifically purified RNAs is limited.

**Figure 6.**
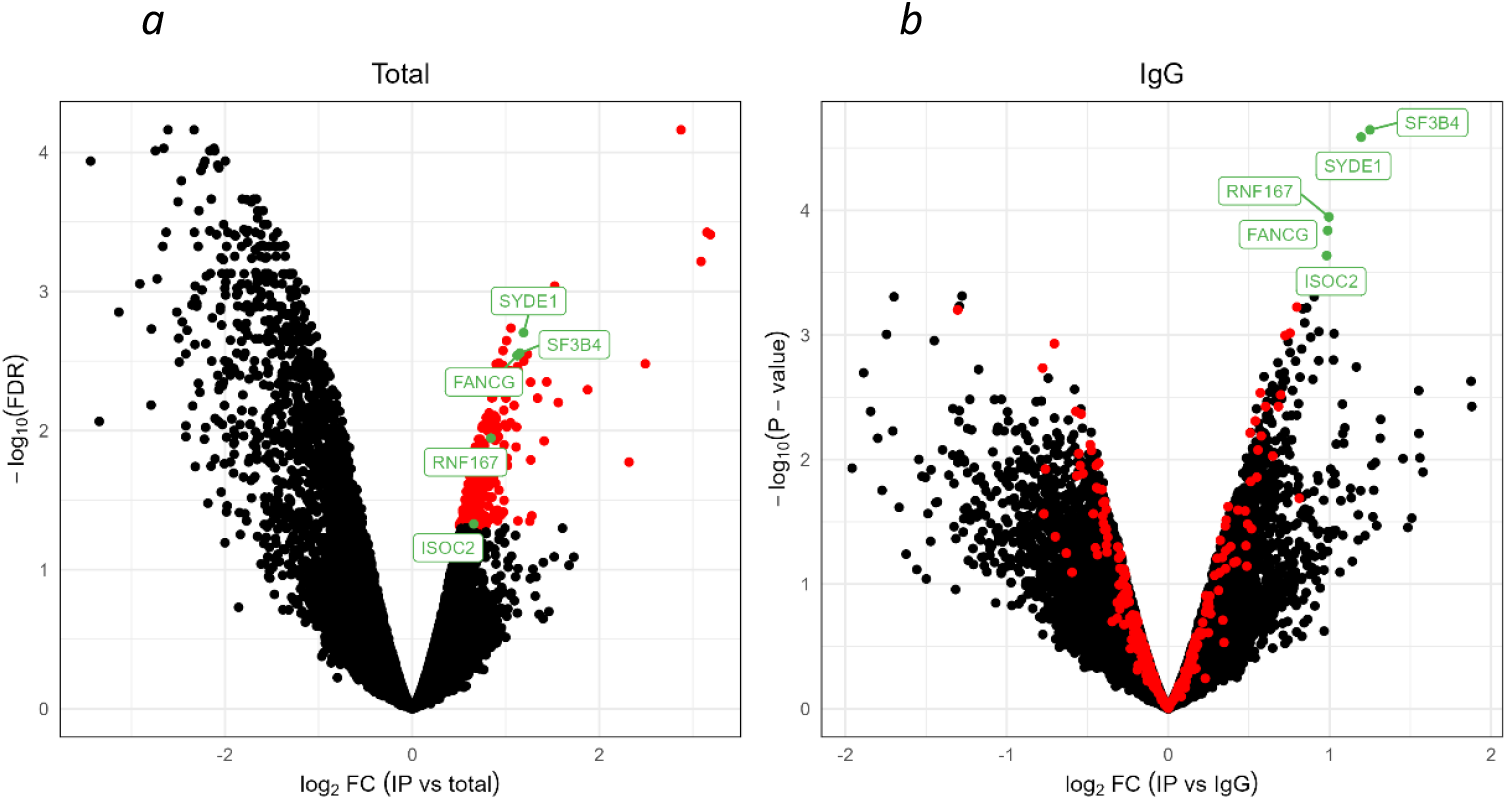
Volcano plots of RNA-Seq changes between IP samples versus total RNA (a) or IgG controls (b). The plots show the log_2_ fold change on the x-axis and the –log_10_ corrected P-value (a) or non-corrected *P*-value (b) on the y-axis. Genes significantly enriched in IP vs total RNA (FDR < 0.05) are highlighted in red. The top most significantly enriched genes in IP vs IgG control are highlighted in green.

We have chosen mRNA SYDE1 coding SYnapse DEfective Rho GTPase Activating Protein 1 for testing NUCB1 mRNA-binding affinity. SYDE1 known as an oncogene in gliomas that can regulate the proliferation and migration of glioma cells (Han *et al*., 2021). According to microRNA target prediction tools (https://www.targetscan.org/vert_80/) in 3′-UTR of SYDE1 mRNA could be target region for miR-27b-3p (Fig. 7). Using SPR was shown that NUCB1 binds SYDE1 with K_D_=314 nM (Table 2).

**Figure 7.**
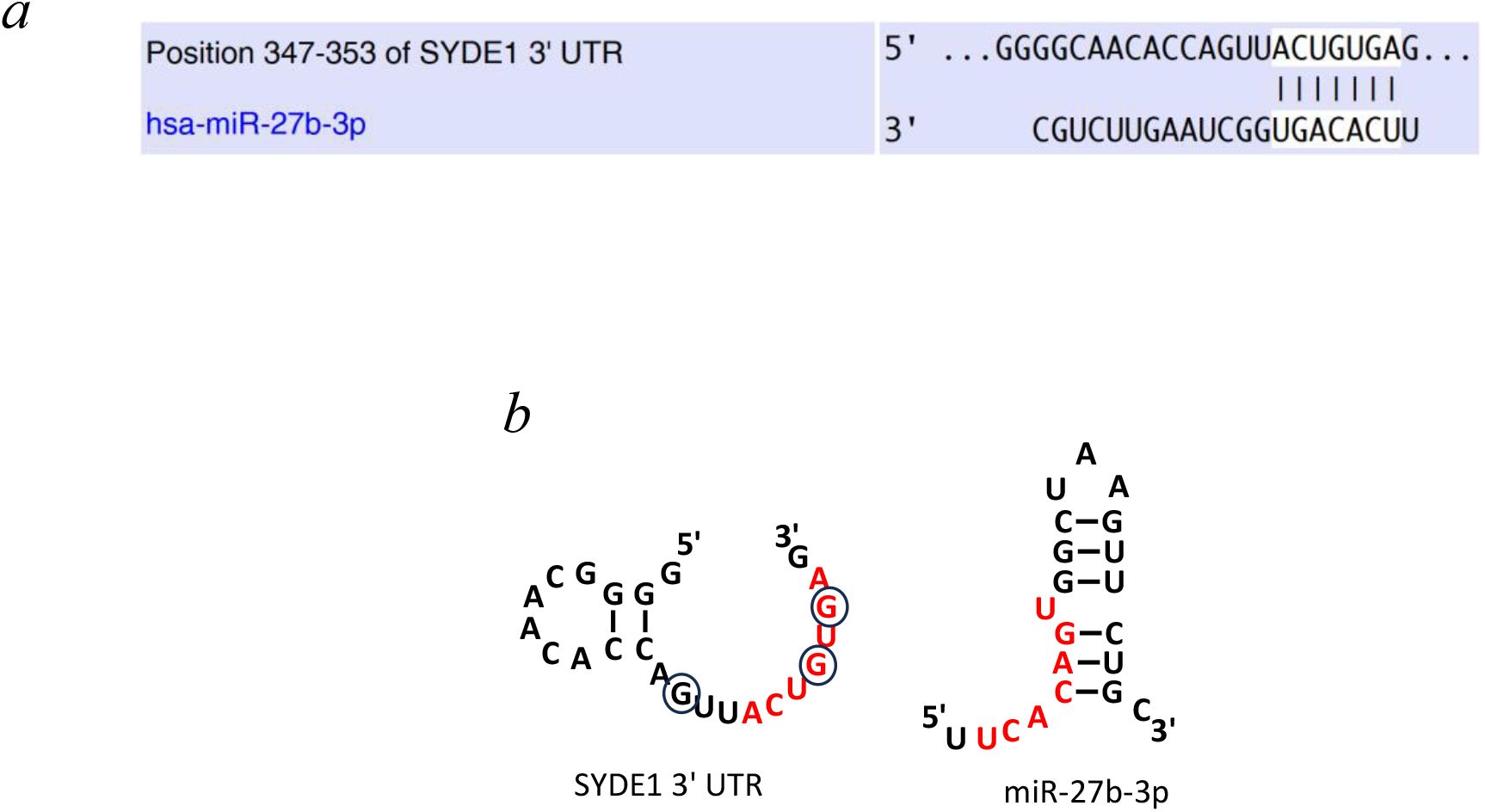
SYDE1 3’ UTR and miR-27b-3p. miR-27b-3p. Alignment of hsa-miR-27b-3p with conserved region 347–353 of SYDE1 3′-UTR using TargetScan (a). Predicted consequential pairing of “seed” region (top) and microRNA (bottom). Secondary structures of SYDE1 3’ UTR and miR-27b-3p (b). The region of 3’UTR SYDE1 that complementary hsa-miR-27b-3p sequence showed as red.

## 3. DISCUSSION

NUCB1 is one of the most abundant Golgi protein with functions related to immunity, apoptosis, calcium homeostasis, G protein signaling, epithelial-to-mesenchymal transition (EMT). NUCB1 bind to E-box DNA (CACGTG) (Sinha *et al*., 2019), interact with necessarily containing guanine ssRNAs and unfold hairpin RNA structures (Mikhaylina *et al*., 2023). NUCB1 was found in endosomes (Larkin *et al*., 2016) and exosomes (Vignesh *et al*., 2021).

In this study we have shown that each of three structural domains of a NUCB1 (DNA-binding region, EF-hands Ca^2+^ - binding region and a leucine zipper) binds RNA fragment with G located in loop. We assumed that the presence of multiple RNA-binding domains would allow the protein to interact with multiple specific RNA regions. Indeed, it was demonstrated using molecular beacon assay that NUCB1 can not only bind but also modulate structures of two RNAs. Moreover, we have shown mRNA-binding activity of NUCB1 *in vivo*.

### 3.1. The potential physiological role of NUCB1 interactions with two RNAs

Our experiments with molecular beacon were similar with experiments with adenine-rich RNA beacon, uridine-rich RNA that can bind with beacon and bacterial/archaeal RNA-chaperone: E.coli Hfq or SmAPs (Lekontseva *et al*., 2020). In Sm protein Hfq and Sm-like archaeal proteins (SmAP) is three different RNA-binding sites (including A-and U-binding site) and these proteins can facilitate the interaction of two complementary RNAs (f.e. mRNA and smallRNA). The addition of RNA-chaperone to the molecular beacon containing A and U nucleotides increased in fluorescence intensity, and the addition in reaction mix second RNA containing U-rich sequence causes sharp increase in fluorescence. The interaction of a chaperone with two RNAs can promote convergence of two partially complementary RNAs increasing the probability of molecular beacon unwinding and resulting in a sharp increase in fluorescence. Thus, it was shown chaperone contribute to the formation of an RNA duplex *in vitro*.

In this study, sharp increases in fluorescence intensity was observed after adding in reaction mix (NUCB1/G-containing loop RNA beacon) second RNA that bind with NUCB1 and contain complementary to beacon region (Fig. 4c, 5c). Apparently, NUCB1 bound to RNA beacon can interact with second RNA and help to form of RNA duplex. In the result, shifting sharp the balance from the hairpin forms of the molecular beacon to the unfolded shape.

It is known many multifunctional RNA-binding eukaryotic proteins according to different functions. One of them is AU-rich RNAs binding protein Hu-antigen R (HuR), containing three RNA-recognized motifs (Ripin *et al*., 2019). Upon stress, HuR binds to the CAT-1 3’UTR in a region positioned downstream of miR-122 sites and causes the repression by displacing the microRNA from mRNA (Bhattacharyya *et al*., 2006). HuR binds both CAT-1 3’UTR and miR-122 (Mukherjee *et al*., 2016).

In assay 2 after adding mir-27b-3p.1 to MB-FOXO the NUCB1 was added (fig.4c, red line). It is clearly visible that the melting of the MB-FOXO is more intense than when NUCB1 added in complex with mir-27b-3p.1. We speculate that NUCB1 can displaced mir-27b-3p.1 from MB-FOXO like in case of HuR protein and mRNA/microRNA. Thus, presumably NUCB1 can not only promote the formation of an RNA duplex, but also destroy it. Naturally, these guesses require further research.

### 3.2. The mRNA-binding activity of NUCB1 in vivo

We conducted mRNA targets of NUCB1 search in HEK-293T cell line using RNA-seq showed in order to confirm RNA-binding activity in vivo. It was found a few mRNA reliable associated with NUCB1 - SYDE1, SF3B4 (Splicing factor 3B subunit 4), FACG (Fanconi Anemia, Complementation Group G) (fig.6). The limited number of target mRNAs may be attributed to several factors: the dependency of RNA-binding ability on post-translational modifications of NUCB1, the requirement for specific RBP complexes, or the expression of certain miRNAs. It’s possible that not all of these factors are fully present in HEK-293T cells and the regulatory role of the NUCB1 may be fully expressed under stressful condition.

We chose SYDE1 mRNA as high statistical significance and the more studied one. Synapse defective 1 (SYDE1), which encodes a Rho GTPase-activating protein is highly expressed in human placenta involving in cytoskeletal remodeling, trophoblast cell migration and acts as an oncogene in glioma (Micale *et al*., 2022; Huang *et al*., 2015; Han *et al*., 2021). SYDE1 is targeted by several microRNA (including miR-200c-3p, miR 27b-3p). Position 347-353 of SYDE1 3’ UTR (target sequence for miR 27b-3p) was selected for investigation of SYDE1 affinity to NUCB1. It was shown that NUCB1 formed complex with SYDE1 fragment (tabl.2). Thus, RNA-seq studies at first time allow confirm mRNA-binding activity of NUCB1 in vivo and identify its potential mRNA targets.

### 3.3. Potential sites of NUCB1 interaction with RNA

Meta-analysis of data from multiple studies using proteomics identified RNA-binding TF including NUCB1 (Oksuz *et al*., 2023). The human genome contains at least 1200 confirmed RBPs (Kelaini *et al*., 2021). Eukaryotic RBP includes heterogeneous nuclear ribonucleoprotein family (hnRNP), the arginine/serine-rich splicing factor protein family (SRSF), RNA-binding motif (RBM) proteins family, proteins with K homology (KH) domain, KH–R3H domains, CSD domain, double-stranded RNA-binding motif (dsRBM) and zinc finger domain (Hentze *et al*., 2025). Most of RBPs contain several RNA-binding domains, including different types of RNA binding motifs. Nevertheless, many of the newly discovered RBPs lack known RNA-binding domains (Hentze *et al*., 2018). Despite its pronounced RNA-binding activity, NUCB1 does not contain any known RNA-binding domains. Nevertheless, there are TFs do not harbor domains characteristic of well-studied RNA binding proteins.

In the work (Sinha *et al*., 2019) pointed out to significant structural similarities of DBD domains of NUCB1 and E-Box binding transcription factor (TF) Myc (a proto-oncogenic protein). The MYC heterodimerizes with MAX protein via its leucine zipper domain and its heterodimer binds to the E-box DNA sequences. Recently was found that Myc binds guanosine-rich RNAs through domains with the arginine-rich motif (Oksuz *et al*., 2023). It was suggested (Li *et al*., 2025) using mutational analysis that two highly conserved Arg-containing sequences (KRR and RQRR), within the basic region of MYC are necessary for its RNA binding.

We found in NUCB1 Arg-containing conserved sequences (RRR, Q(E)RR(K), RR(K)) and marked them on the NUCB1 dimer structure predicted by the Alphafold3 (fig. 8).

**Figure 8.**
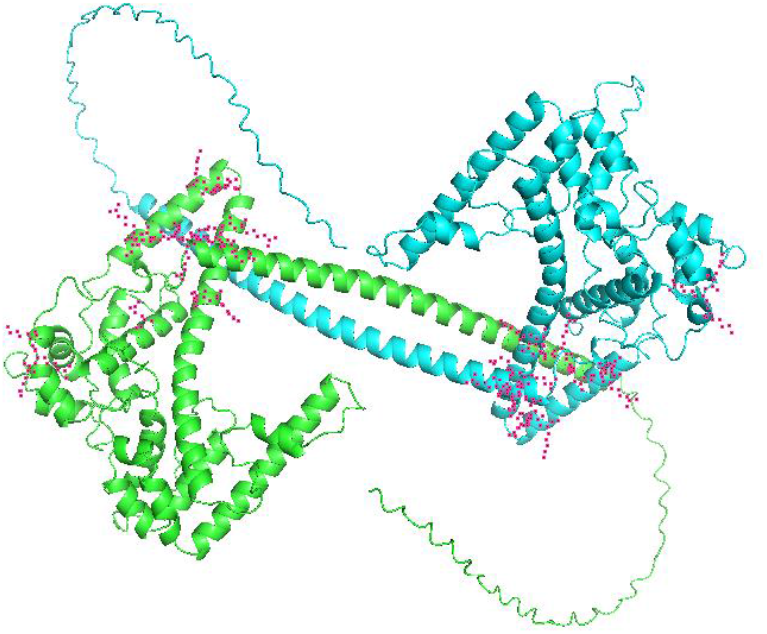
Structural model of dimer NUCB1 predicted by the AlphaFold3. Monomers are shown in blue and green, the side groups of residues (RRR, Q(E)RR(K), RR(K), KK) are shown in red. C-terminals (411-461 aa) including polyQ sequences are unordered.

Arginine “patches” was found in DNA-binding, Ca^2+^ - binding and a leucine zipper domains and can form two groups on the surface of each monomer. At present time the structure of the calcium-binding domain of the NUCB1 (pdb id 1SNL, de Alba *et al*., 2004) is known only. The structure of full protein is unknown. Therefore, it is difficult to say which amino acid residues and structural elements of the protein are involved in the RNA-protein interaction certainly. However, since only the full-length NUCB1 has RNA chaperone activity (Mikhaylina *et al*, 2023), obviously it is necessary for the realization of this activity to have all the RNA-binding sites of the NUCB1 dimer. It should also be noted that not all three RNA-binding sites of the protein could be structural available for interaction with the RNAs in the NUCB1 dimer structure.

## 4. Conclusion

Our findings on the RNA-binding and RNA chaperone activity of NUCB1 *in vitro* and first RNA-seq experiment *in vivo* allow making several assumptions about its influence on translation regulation. Presumably, binding of NUCB1 to G-containing single strand sites of the 3’ UTR of mRNA and microRNA can facilitate RNA-RNA interaction (annealing) or displace microRNA from mRNA. Besides, interaction of NUCB1 with mRNA and/or microRNA can shield them from RNases. NUCB1 can also modulate the structure of mRNA by binding to the G-containing single-stranded regions of mRNA.

## 5. Materials and Methods

### 5.1. Production and purification of NUCB1 and its truncated forms

NUCB1 protein was obtained as previously described (Mikhaylina *et al*., 2023).

Genes of the truncated forms NUCB1: NUCB1 DBD and NUCB1 LZ, were PCR-amplified using primers For NUCB1 (EcoRI) and Rev NUCB1 DBD (HindIII); For NUCB1 LZ (EcoRI) and Rev NUCB1 (HindIII), respectively (Table 3), and plasmid pET-28a/NUCB1 as a template. The genes were cloned in the pET-28a vector similarly to NUCB1 gene. The primers contain sites for cleavage by site-specific restriction endonucleases EcoRI and HindIII necessary for inserting the gene into pET-28a vector. The obtained genetic constructions were validated by sequencing. NUCB1 DBD contains 31-219 amino acid residues of NUCB1, and NUCB1 LZ contains 327-461 amino acid residues of NUCB1.

**Table 3.**
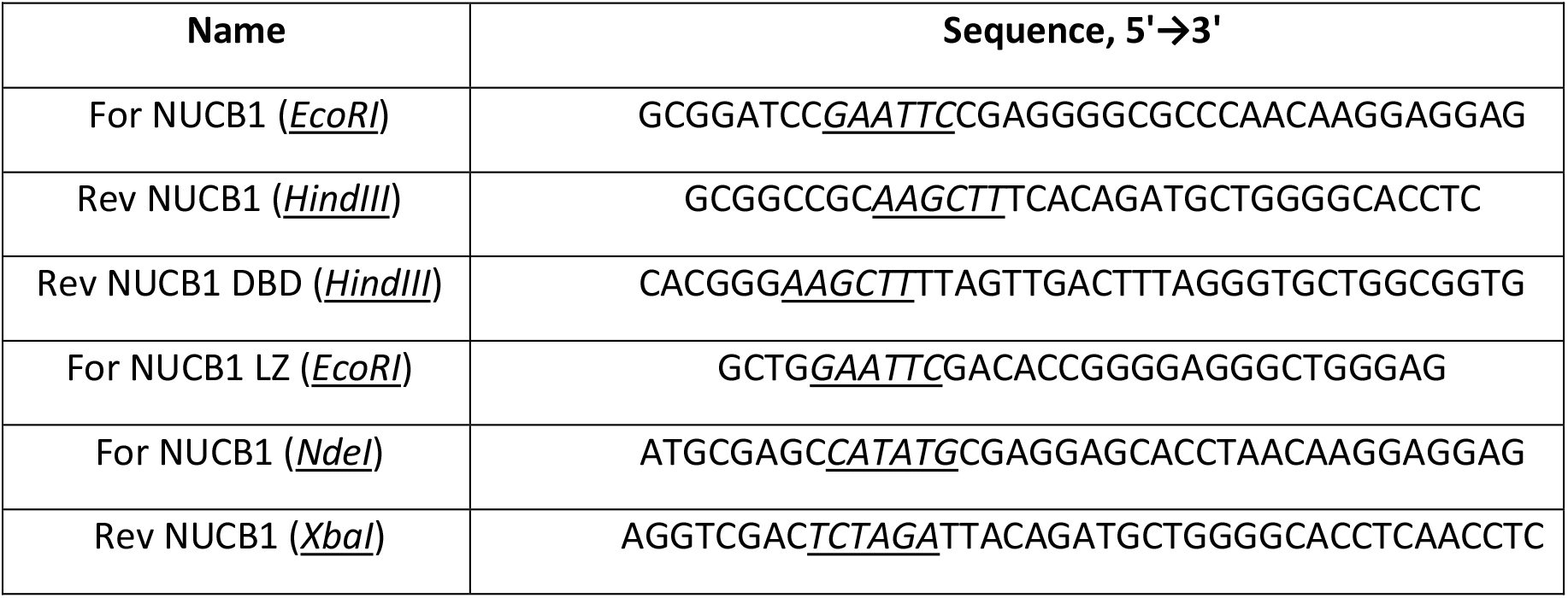
Primers used in this study.

Protein purification was carried out according to the previously described method (Mikhaylina *et al*., 2023).

### 5.2. Molecular beacon melting assay

A Cary Eclipse fluorescence spectrometer (Varian, USA) was used in the assay. All measurements were performed in a 0.3×0.3 cm cuvette at a constant temperature of 22°C; the excitation wavelength was 496 nm, and emission wavelength was 519 nm.

#### 5.2.1. RNA melting properties of NUCB1

To study the RNA chaperone properties of NUCB1, chemically synthesized molecular beacon-FOXO1 3’-UTR (MB-FOXO) and molecular beacon-microRNA-27b-3p (MB-mir) were used (Syntol, Russia):

MB-FOXO 5’-FAM-***GGUCC***GUAAUACUGUGAU***GGACC***-RTQ-3’,

MB-mir: 5’-FAM-***GGUCC***UUCACAGUGGCUAAGUUCUGCCC***GGACC***-RTQ-3’

A MB-mir is a hairpin RNA (highlighted in bold italics in the sequences) containing the RNA sequence in a loop with a FAM fluorophore at one end and an RTQ1 quencher (analog of BHQ1) at the other end (Figure 3a, 5a). In the initial state, the 5’-and 3’-ends of MB-miR are in close proximity, so the fluorescence is quenched.

Sequence of miR-27b-3p.1 RNA contains, in addition to the mir-27b-3p sequence, a beacon helix sequence (bold italic) to form a single-stranded NUCB1 binding site, as it is in MB-mir.

Analysis of protein-induced changes in fluorescence intensity was performed for NUCB1 as it was described in our previous work (Mikhaylina *et al*., 2023).

NUCB1 was added to MB-RNA in molar ratio 10:1 (2mkM:200 nM) in 10 mM Tris-HCl (pH 8.0) and 100 mM NaCl (buffer A). The volume of reaction mix was 100 µl. Three independent melting curves were obtained for each MB-mir-protein combination.

#### 5.2.2. RNA beacon melting experiments in presence of second RNA

For competition experiments we used chemically synthesized FOXO1 contained complementary site to MB-mir; miR-27b-3p, and miR-27b-3p.1 contained complementary site to MB-FOXO and oligoC as negative control (Syntol, Russia):

FOXO1: 5’-GUAAUACUGUGAUGGGCC-3’,

miR-27b-3p: 5’-UUCACAGUGGCUAAGUUCUGC-3’,

miR-27b-3p.1: 5’-***GGUCC***UUCACAGUGGCUAAGUUCUGCCC***GGACC***-3’,

oligo(C): 5’-GUGGUCAGUCGAGUGG-CCCCCC-3’.

The miR-27b-3p.1 sequence differs from the miR-27b-3p by the regions that form RNA hairpin (highlighted in bold italics in the sequences), which ensures the formation a single-stranded NUCB1 binding site in loop (Fig.5a). We used the buffer A for all RNA beacon-melting experiments, volume of reaction mix was 100 µl.

To assess effect of complementary RNAs (mir-27b-3p, mir-27b-3p.1, FOXO) on MB-RNAs (MB-FOXO, MB-mir) melting by NUCB1 the complementary RNA was added at 20:1 molar ratio to MB-RNA (4 mkM:200 nM). NUCB1 was added at 10:1 molar ratio to MB-RNA (2 mkM:200 nM, respectively).

In another version of experiment mix of NUCB1 and miR-27b-3p (Fig. 3b) or mix of NUCB1 and miR-27b-3p.1 (Fig.4c) in molar ratio 1:2 (2 mkM:4 mkM, respectively) were added to the 200 nM MB-FOXO.

For the control experiments water, buffer A or oligo(C) RNA were added to the MB-RNA/NUCB1 mix instead of complementary RNA.

It was shown, the time dependence of the effect of adding partially complementary RNA to the beacon, with which the NUCB1 does not bind (Fig.3c). In reaction mix (MB-FOXO/NUCB1) 4 mkM of miR-27b-3p was added at equal intervals: 10 min, 20 min, 30 min and 40 min. In the control experiment, water was added to MB-FOXO after 10 min after NUCB1 addition.

Three independent melting curves were obtained for each MB-miR-protein combination.

### 5.3. Analysis of NUCB1-RNA interaction

Kinetic analysis of NUCB1 and its truncated forms interaction with specific RNA fragments was performed by surface plasmon resonance technique (Mikhaylina *et al*., 2014) using the ProteOn XPR36 system (Bio-Rad, USA).

The RNA-binding properties of NUCB1 and its truncated forms were tested using 5’-biotinylated chemically synthesized RNA oligomers (Syntol, Russia) listed below:

miR-200a-3p: 5’-GUACCGAGCUCGAAUUUAACACUGUCUGGUAACGAUGU-3’

FOXO1: 5’-GUACCGAGCUCGAAGUAAUACUGUGAUGGGCC-3’,

mir-27b-3p: 5’-GUACCGAGCUCGAAUUCACAGUGGCUAAGUUCUGC-3’,

miR-27b-3p.1 5’-GUACCGAGCUCGAA***GGUCC***UUCACAGUGGCUAAGUUCUGCCC***GGACC***-3’

SYDE1: 5’-GUACCGAGCUCGAAGGGGCAACACCAGUUACUGUG-3’

Fourteen nucleotides (underlined in sequences) at the 5’-end of oligo RNAs were added to increase the accessibility of short RNA fragments for interaction with proteins. Biotinylated RNA fragments were applied onto the NLC sensor chips with immobilized avidin (Bio-Rad, United States).

Five different concentrations of the analyte samples (NUCB1 or truncated forms) were prepared by serial dilution in a solution containing 50 mM Tris-HCl (pH 8.0), 200 mM NaCl, and 0.1 % Tween-20 (TNT) for each set of sensorgrams. The samples were injected at a flow rate of 30 µl/min. The injection step included a 300 s association phase followed by a 2000–6000 s dissociation phase in TNT buffer. All binding experiments were performed at 25°C. Each immobilization strategy and kinetic analysis was repeated at least three times.

Kinetic analysis was performed by globally fitting curves describing the simple 1:1 bimolecular model to the set of three to five sensorgrams using BIAEvaluation software.

### 5.4. RNA immunoprecipitation and sequencing

#### 5.4.1. Polyclonal antibodies specific to NUCB1

Rabbit serum containing antibodies specific to the NUCB1 protein was obtained at the Laboratory of Immunochemistry, Pushchino Branch, Shemyakin–Ovchinnikov Institute of Bioorganic Chemistry, Russian Academy of Sciences, Russia. Rabbit was immunized with recombinant NUCB1 protein, obtained as described previously (Mikhaylina *et al*., 2023).

#### 5.4.2. Cell cultivation and transfection

Gene of the NUCB1 was PCR-amplified using primers For NUCB1 (NdeI) and Rev NUCB1 (XbaI) (Table 3) and plasmid pET-28a/NUCB1 as a template. The primers contain sites for cleavage by site-specific restriction endonucleases NdeI and XbaI necessary for inserting the gene into pcDNA3-HA vector. The obtained genetic constructions were validated by sequencing.

HEK293T cells (originally obtained from ATCC) were cultivated in Dulbecco’s Modified Eagle’s Medium (DMEM) supplemented with 10% fetal calf serum, 2 mM glutamine, 100 U/ml penicillin, and 100 µg/ml streptomycin. The cells were incubated at 37°C in a humidified atmosphere containing 5% CO_2_ and passaged by standard methods.

To obtain NUCB1-overexpressing cells, the cells were cultured on a 6-well plate to a density of 80-90%. The cells were transfected using Lipofectamine 3000 (Invitrogen). For transfection, 5 μg of the plasmid pcDNA3-HA-NUCB1 or 5 μg of pcDNA3-HA (as control) was incubated with 10 μl of the P3000 reagent and 7.5 μl of the Lipofectamine 3000 in 250 μl Opti-MEM for 5 min and then added to the growth medium. 12 hours later, the cells were plated onto a new 100-mm dish and cultivated for 36 h.

#### 5.4.3. RNA immunoprecipitation

RNA immunoprecipitation was performed as described by Tenenbaum *et al*., (Tenenbaum at al., 2000) with some modifications. Briefly, cells were washed twice with ice-cold PBS and removed from culture plates with a scraper. Then cells were resuspended in approximately two pellet volumes of lysis buffer containing 10 mM Hepes-KOH, pH 7.0, 100 mM KCl, 5 mM MgCl_2_, 0.5% Nonidet P-40 with 1 mM DTT, 100 U/ml RNase Inhibitor (Thermo Fisher Scientific), 0.2% vanadylribonucleoside complex (Fluka), and Protein Inhibitor Cocktail (Roche) added fresh at the time of use. The lysed cells were then frozen and stored at ™80°C. At the time of use, the cell lysate was thawed and centrifuged at 16,000 g in a microfuge for 10 min at 4°C. For immunoprecipitation, Protein G Sepharose beads (GE Healthcare) were swollen 1:5 V/V in NT2 Buffer (50 mM Tris-HCl, pH 7.6, 150 mM NaCl, 1 mM MgCl_2_, 0.05% Nonidet P-40) supplemented with 5% BSA. A 300 μl aliquot of the 1:5 V/V pre-swollen protein G bead slurry was used per immunoprecipitation reaction and incubated overnight at 4°C with an excess of antiNUCB1 (rabbit antiserum containing antibodies specific to the NUCB1) or control antibodies (unimmunized rabbit serum) (20 μg). The antibody-coated beads were washed with ice-cold NT2 buffer and resuspended in 900 μl of NT2 buffer supplemented with 100 U/ml RNase Inhibitor, 0.2% vanadylribonucleoside complex, 1 mM DTT, 20 mM EDTA. The beads were briefly vortexed, and 100 μl of cell lysate was added and immediately centrifuged, and a 100 μl aliquot removed to represent total cellular RNA. The immunoprecipitation reactions were performed at room temperature for 2 h and then beads were washed 4 times with ice-cold NT2 buffer followed by 2 washes with NT2 buffer supplemented with 1 M urea. Washed beads were resuspended in 100 μl NT2 buffer.

#### 5.4.4. Library preparation and sequencing

RNA from immunoprecipitated samples (NUCB1 or IgG antibodies) and total RNA was isolated using QIAzol (Qiagen) and Direct-Zol RNA Microprep kit (Zymo Research). Sequencing libraries were prepared with NEBNext Ultra Directional Kit (NEB), and mRNA was enriched with NEBNext Poly(A) Module (NEB). All steps followed the manufacturers’ protocols. Libraries were sequenced on Illumina Novaseq 6000 (Skoltech Genomics Core Facility).

#### 5.4.5. Data Processing

FASTQ files for 6 samples (2 replicates per experiment type: total RNA, NUCB1 IP, and IgG IP), each containing 31-39 million paired-end reads, were preprocessed with cutadapt (v.3.5) (Marcel, 2011) *--max-n 0.1 --discard-trimmed -A AGATCGGAAGAG* to remove fragments with short insert size or large fraction of ‘N’ bases, leaving from 81 to 91% of the total read pairs. The next step included mapping to the GRCh38 human genome with STAR (v.2.7.10) (Dobin *et al*., 2013) *-- quantMode GeneCounts --alignSJDBoverhangMin 1* using GENCODE (v.38) comprehensive annotation. Gene counts from STAR corresponding to the 2nd read strand aligned with RNA were used in further analysis. Differential expression analysis was performed in R using edgeR v4 (Baldoni *et al*., 2024). Genes with counts per million (CPM) > 0.5 in at least 4 samples (∼10 reads) were normalized (TMM), with dispersion estimated (*estimateDisp*). Differential expression (IP vs total RNA or IgG) was tested using a generalized linear model (*glmQLFit, glmQLFTest*). *P*-values were corrected by FDR (Benjamini-Hochberg).

